# A novel multi-functional role for the MCM8/9 helicase complex in maintaining fork integrity during replication stress

**DOI:** 10.1101/2021.12.08.471807

**Authors:** Wezley C. Griffin, David R. McKinzey, Kathleen N. Klinzing, Rithvik Baratam, Michael A. Trakselis

## Abstract

The minichromosome maintenance (MCM) 8/9 helicase is a AAA^+^ complex involved in DNA replication-associated repair. Despite high sequence homology to the MCM2-7 helicase, an active role for MCM8/9 has remained elusive. We interrogated fork progression in cells lacking MCM8 or 9 and find there is a functional partitioning. Loss of MCM8 or 9 slows overall replication speed and increases markers of genomic damage and fork instability, further compounded upon treatment with hydroxyurea. MCM8/9 acts upstream and antagonizes the recruitment of BRCA1 in nontreated conditions. However, upon treatment with fork stalling agents, MCM9 recruits Rad51 to protect and remodel persistently stalled forks. The helicase function of MCM8/9 aids in normal replication fork progression, but upon excessive stalling, MCM8/9 directs additional stabilizers to protect forks from degradation. This evidence defines novel multifunctional roles for MCM8/9 in promoting normal replication fork progression and promoting genome integrity following stress.

## Introduction

Accurate genomic duplication during S-phase is vital such that each daughter cell is guaranteed a copy of the complete, unadulterated genome. Several thousand replication complexes are licensed and fired with temporal and spatial precision to ensure ephemeral but complete DNA replication.^1,2^ A variety of challenges, including DNA template damage, DNA secondary structure, or DNA-protein blocks, are often encountered by the replication machinery.^3,4^ These challenges often stall replication forks, either temporarily or more persistently, and if not rescued or restarted by a variety of DNA damage responses (DDR) can collapse into DNA double-strand breaks (DSBs). Such breaks are hallmarks of chromosomal instability and can contribute to cancer development, aging, and infertility.^5–7^

Fortunately, cells have evolved several failsafe mechanisms to thwart the deleterious outcomes of replication fork stalling and collapse through activation of fork protection pathways.^8^ These pathways are coordinated by the ATR kinase, which signals a variety of downstream stress responses that inhibit cell cycle progression, suppress late origin firing, and ensure stabilization and recovery of stalled or reversed replication forks.^9^ Replication fork reversal is one mechanism that continues to gain support as a general defense to protect stressed forks and prevent fork collapse.^10–12^ The fork reversal/restart mechanism can be sub-divided into three basic steps: 1) SNF2 enzyme-mediated annealing of newly synthesized and parental DNA strands to form a regressed arm and a four-way DNA junction or ‘chicken foot’ structure, 2) removal of damage or replication block, and 3) re-installation and restart of the replication complex. Additionally, many proteins important for homologous recombination (HR) repair of DSBs (*e.g*. BRCA1/2, Rad51, Mre11, etc.) moonlight at stalled or reversed forks to prevent genomic instability through fork protection/restart or recombination.^13^

The minichromosome maintenance (MCM) 8 and 9 are ATPases associated with a variety of cellular activities (AAA^+^) and are homologs within the MCM family of proteins. While MCM2-7 forms the core of the replicative helicase, MCM8 and 9 form a discrete hexameric helicase complex implicated in HR-mediate repair. Studies have linked the loss of MCM8 or 9 to primary ovarian failure (POF), infertility^14–16^, and cancer^17^, with more than 400 different mutations in both MCM8 and 9 cataloged in genome databases.^18^ Many of these mutations lack sufficient characterization detailing the molecular and cellular effects that contribute to disease initiation and progression, and so, the importance of MCM8 and 9 in maintaining genomic stability is still being deciphered. Many of these reports show a direct link between a functional MCM8/9 complex and successful meiotic or mitotic HR. Indeed, both mice and humans with non-functional MCM8/9 display reproductive system abnormalities including infertility, sex-specific tumor formation, sensitivity to DNA damaging agents, and defects in HR processing.^19,20^ Furthermore, loss of MCM8/9 impairs HR-mediated fork rescue due to decreased recruitment of the MRN helicase/nuclease complex, Rad51 recombinase, and RPA single-stranded (ss-) DNA binding protein after cisplatin (Cis-Pt) treatment.^21^

Despite a high sequence homology to MCM2-7, a precise function of MCM8/9 at the DNA replication fork has remained enigmatic. Early reports debated whether MCM8/9 was required for prereplication complex (preRC) assembly^22^ or only utilized at functionally activated replication forks but not required for replication.^23–26^ Although more recent studies have focused primarily on MCM8/9’s participation in HR, there were several lines of evidence hinting that MCM8/9 may play an active role during DNA replication. For example, replication forks from cells lacking MCM8 or 9 stalled or collapsed nearly 2-fold more than controls cells suggesting loss of MCM8/9 sensitizes forks to replication stress.^19^ Additionally, when MCM2 is rapidly degraded, MCM8/9 can fill in and allow for DNA dependent synthesis, albeit at a significantly slower overall rate.^27^ Furthermore, iPond supports the presence of MCM8/9 at active replisomes at a similar level to other *bona fide* replication proteins.^28^ Together, the bulk of the evidence suggests that MCM8/9 actively contributes to genomic integrity by directly promoting replisome progression through the stabilization and protection of DNA replication forks during active elongation.

Here, we report a novel role for the MCM8/9 complex in maintaining replication fork stability during progression, stalling, and reversal. By integrating single-molecule DNA fiber and neutral comet assays along with flow cytometry and immunofluorescence analyses, we show that MCM8/9 knockout (KO) cell lines exhibit reduced rates of DNA synthesis, delayed cell cycle progression, and increased markers of genomic instability as a consequence of reduced replication fork stability. Collectively, our data support a multi-functional model whereby the helicase domain of MCM8/9 antagonizes BRCA1 dependent fork reversal, stabilization, and processing during normal fork progression. However, upon excessive stalling, MCM8/9 recruits Rad51 through a BRCv motif in the C-terminal extension of MCM9 to protect and reverse stalled forks. Altogether, these results confirm a direct role for MCM8/9 in maintaining genomic integrity by stabilizing the active replication fork.

## Results

### Knockout of the MCM8/9 complex slows DNA synthesis progression and sensitizes cells to replication-induced DNA damage

Recently, multiple proteins with established roles in HR repair have documented activity in maintaining replication fork progression and integrity.^29^ Since MCM8/9 are homologous to the MCM2-7 replicative helicase complex and are involved in HR repair, we hypothesized that MCM8/9 may also be involved in actively maintaining fork integrity during replication. Previous observations have suggested that cells lacking MCM8 or 9 exhibit reduced growth rates^19,27^, which could be explained by compromised fork stability. To assess this possibility, we measured DNA fiber lengths for 8^KO^ or 9^KO^ compared to WT cells to monitor fork progression at different times in the absence of exogenous damage (**Fig. 1a-d**). Mean CldU track length values were then plotted as a function of time and fit to a simple linear regression to obtain apparent replication rates (**Fig. 1e**). The CldU track length in wild-type (WT) 293T cells increased at a rate of ~0.25 μm per minute, which corresponds to a DNA synthesis rate of 12.2 base pairs per second (**Fig. 1b** and **1e**, black circles). As expected, both 8^KO^ and 9^KO^ cells exhibited 3-4 fold reduced CldU track length rates of 0.07 and 0.09 μm per minute, which correspond to DNA synthesis rates of 3.4 and 4.4 base pairs per second, respectively (**Fig. 1c-d**, and **1e**, blue open circles and red open squares, respectively).

**Figure 1.**
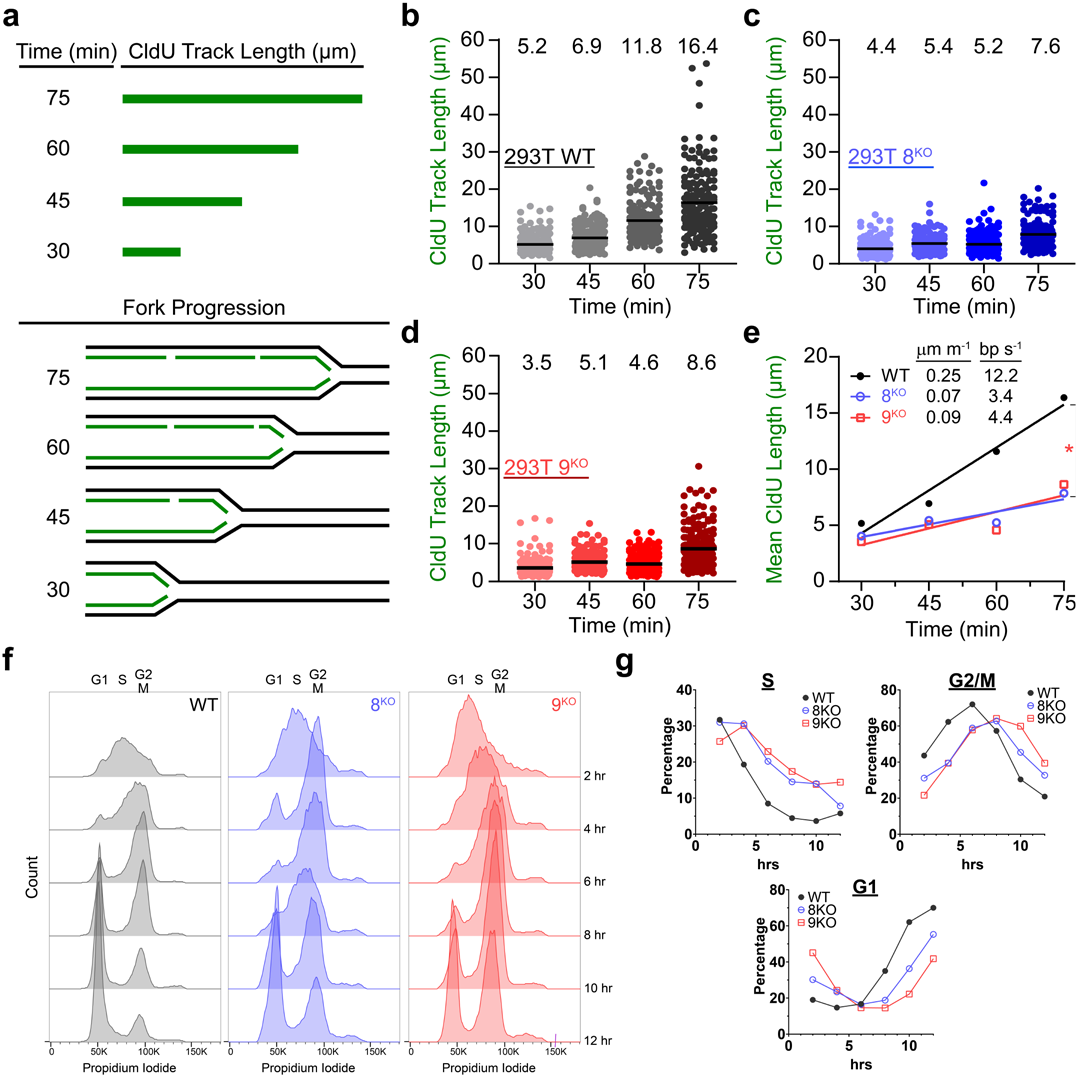
DNA replication rates are reduced in the absence of MCM8/9. **a** Schematic of replication fork progression assay. Cells were treated with 50 μM CldU for the indicated time intervals and the CldU track lengths (upper panel) were measured as a readout of replication progression (bottom panel). CldU track lengths of **b** 293T WT, **c** 8^KO^, or **d** 9^KO^ cells. CldU lengths were measured with ImageJ software and the corresponding mean value of each time point are indicated at the top of the graph. **e** Mean CldU track length values from **b-d** were plotted as a function of time to obtain apparent replication rates. WT (black circles, ●), 8^KO^ (blue circles, 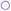), 9^KO^ (red circles, 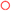). P-values indicate significant differences in the slope (*P< 0.05). **f** Cells were synchronized with a double thymidine block, released into S-phase, and then the chromosome content was monitored by FACS. **g** After gating and quantification of S, G2/M, and G1 populations, the percentages were plotted as a function of time.

Both 8^KO^ or 9^KO^ cells grew in culture at a qualitatively observable slower rate than WT cells. To directly quantify S-phase progression, we performed cell synchronization experiments with a double thymidine block at the G1-S phase boundary. After release, cell cycle progression through S-phase using was monitored by fluorescence activated cell sorting (FACS) (**Fig. 1f**), gating cell populations in S, G2/M, and then G1 by propidium iodide signal (**Fig. 1g**). There is a 2-3 hour delay through S-phase for 8^KO^ or 9^KO^ cells compared to wild-type that translates to an overall delay in cell division (G2/M) and continues through the next G1 phase. This provides strong evidence that the MCM8/9 complex aids in normal replication fork progression and that the loss of MCM8/9 likely results in reduced replication fork stability and genome instability.

Previously, several laboratories including ours have shown that MCM8/9 form nuclear foci, upon damage, primarily from DNA crosslinking agents or after direct DSBs induced from ionizing radiation.^24,30,31^ This implicates MCM8/9 in HR, but there is limited information investigating a possible role during fork progression/stalling. To directly examine whether MCM8/9 are involved in maintaining genomic instability during replication stress, a C-terminal GFP-tagged MCM9 fusion construct was transfected into WT 293T cells, after which cells were treated with 2 mM hydroxyurea (HU) for 4 hours, and MCM9 foci formation was monitored (**Fig. 2a**). Cells treated with HU exhibited more MCM9-dependent foci compared to GFP alone and nontreated controls (**Fig. 2a**, white arrows), indicating that the MCM8/9 complex responds to mild stressors that induce replication fork stalling.

**Figure 2.**
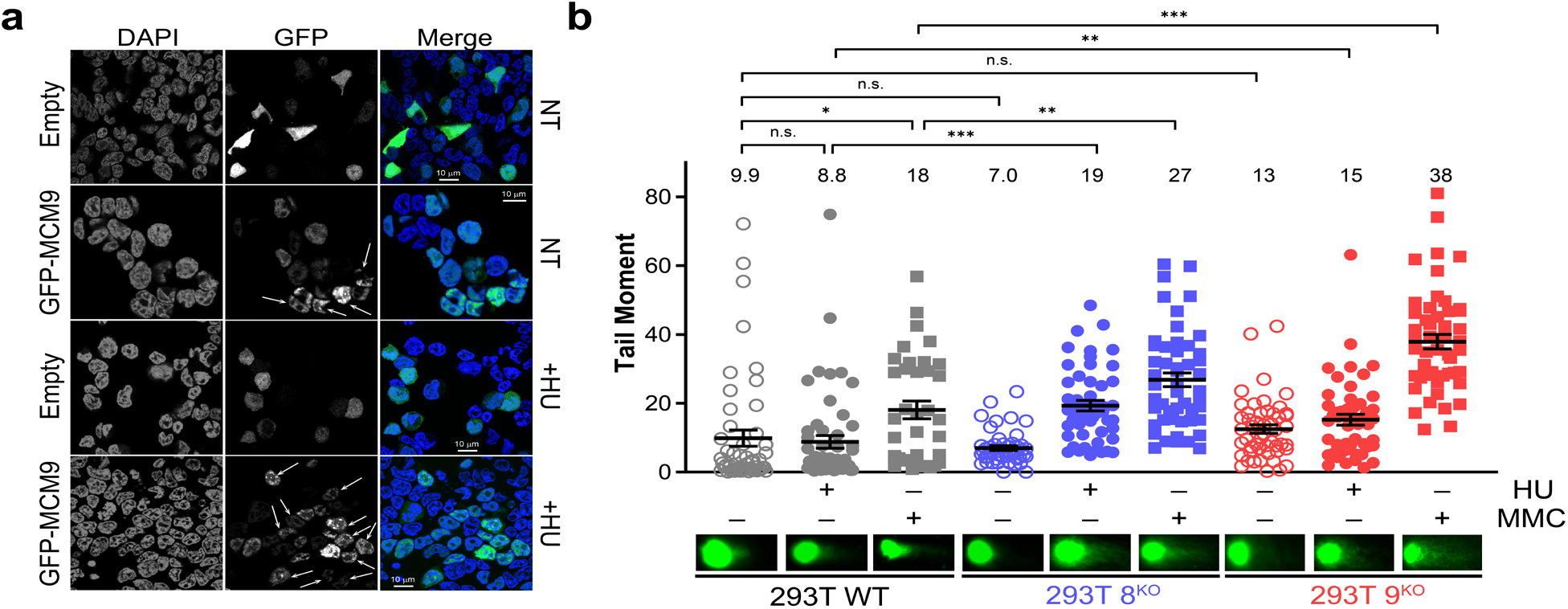
MCM8/9 protect cells from replication-induced DNA damage. **a** 293T cells were transfected with GFP-MCM9 fusion construct or GFP alone, nontreated (NT) or treated with 2 mM HU for 4 hours (+HU), and then imaged by confocal microscopy. White arrows indicate distinct nuclear GFP-MCM9 foci. Scale bars represent 10 μm. **b** Quantification of the tail moment in a neutral comet assay for 293T WT (grey), 8^KO^ (blue), or 9^KO^ (red) cells (○) treated with 2 mM HU for 4 hours (●) or 3 μM MMC for 6 hours (■) A Mann-Whitney test was used to calculate P-values (n.s. nonsignificant, *P< 0.05, **P< 0.01, ***P< 0.001). Representative comets are shown below the graph for each condition.

Replication-associated DNA damage in both 8^KO^ and 9^KO^ cells was directly measured and compared to WT cells using a neutral comet assay to detect DSBs (**Fig. 2b**). Upon treatment of 8^KO^ and 9^KO^ with 2 mM HU for 4 hours, there was a statistically significant (~2.2 and ~1.7-fold, respectively) increase in tail moment values compared to WT cells. Addition of mitomycin C (MMC) to 8^KO^ or 9^KO^ cells showed a similar trend with a ~1.5- and ~2.1-fold increase in tail moment compared to WT, respectively. To show specificity, comet tail moments are rescued by the expression of untagged MCM9 from an IRES2 plasmid after treatment with HU (1.5-fold reduction) or MMC (1.9-fold reduction) (**Supplementary Fig. 1**).

To further investigate the prevalence of DNA breaks occurring in 8^KO^ or 9^KO^ cells, γH2A.X foci were probed in nontreated or HU treated cells (**Fig. 3a-b**). γH2A.X foci are surrogate markers of DNA damage and early effectors of the DSB repair pathway.^32,33^ Interestingly, both 8^KO^ or 9^KO^ cells showed significant increases in γH2A.X foci in nontreated cells consistent with the hypothesis that loss of MCM8/9 results in defective replication that induces genomic stress (**Fig. 3c**). Nontreated WT cells were essentially void of any γH2A.X foci. This effect was enhanced overall with HU treatment, where significantly more foci were again found in 8^KO^ or 9^KO^ cells compared to WT cells (**Fig. 3d**). These results indicate that cells lacking MCM8/9 are more susceptible to DNA damage-inducing events, likely initiated by reduced fork stability during replication.

**Figure 3.**
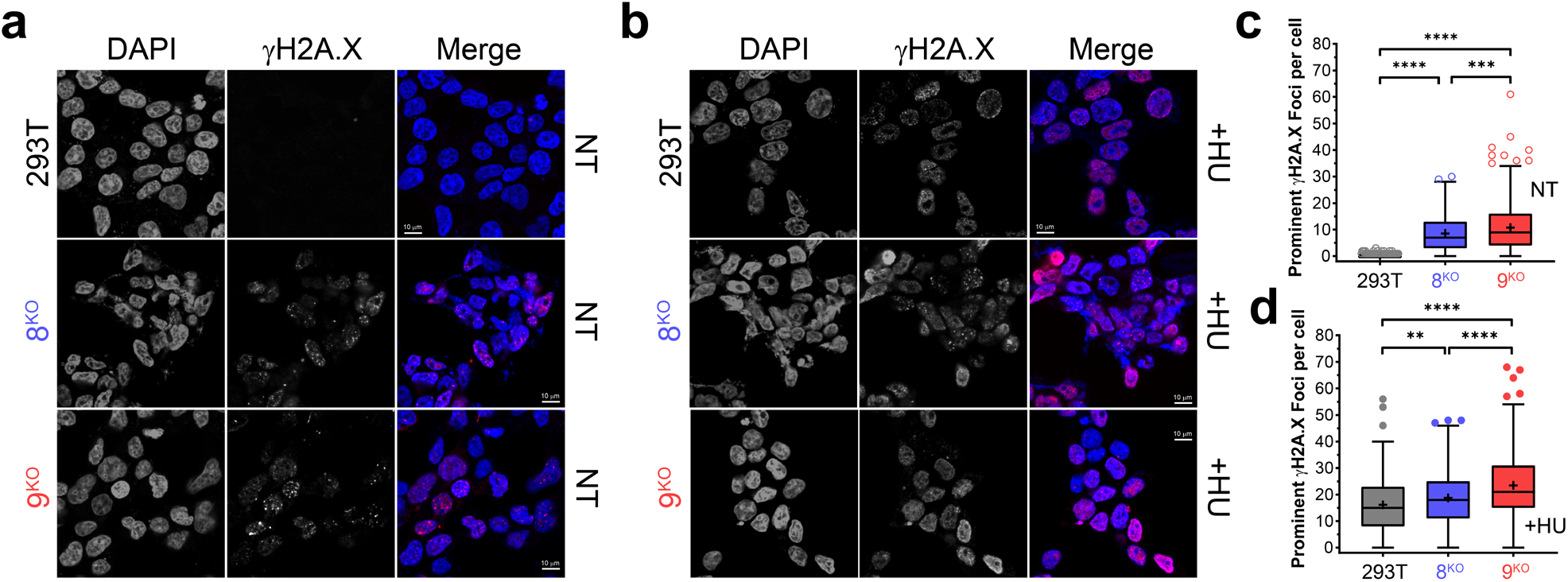
MCM8/9 protects cells from breaks during normal replication and stalling. 293T (grey), 8KO (blue), or 9KO (red) were examined for either **a** native H2A.X foci in nontreated (NT) cells or **b** after treatment with 2 mM HU for 4 hours (+HU) and then imaged by confocal microscopy. Prominent H2A.X foci were quantified per cell using Image J and plotted as a Tukey box and whiskers for **c** NT or **d** HU treated cells with + representing the mean. (**P< 0.01, ***P< 0.001, ****P<0.0001).

### MCM8/9 maintain replication fork integrity during stress and reversal

As MCM8/9 appear to be involved in maintaining replication fork integrity both transiently (during normal synthesis) and during prolonged stress (as shown with HU), we hypothesized that MCM8/9 may act in a similar manner as other HR proteins to stabilize stalled or reversed replication forks.^34,35^ To examine this possibility, we measured replication fork stability in WT, 8^KO^, and 9^KO^ cells by DNA fiber analysis (**Fig. 4**). Interestingly, under normal conditions, both 8^KO^ and 9^KO^ cells exhibit a smaller, statistically significant reduction in the mean IdU/CldU ratio value compared to WT (**Fig. 4a**, compare plots with open circles), suggesting a defect in replication fork stability upon loss of the MCM8/9 complex. This reduction in mean IdU/CldU values was more pronounced in the presence of 2 mM HU, which stimulates more persistent replication stress and initiates fork reversal (**Fig. 4a**, compare lanes with filled circles), indicating that loss of MCM8/9 further sensitizes replication forks to degradation following stress. Furthermore, transfection of WT MCM8 or MCM9 constructs into the respective KO cells partially restores replication fork stability following 2 mM HU treatment, as indicated by the increase in mean IdU/CldU value compared to GFP alone transfected controls (**Fig. 4b**, compare plots with filled circles). We note that DNA fiber measurements and quantifications for transfection of GFP into 9^KO^ cells (**Fig. 4b**, plots 5&6) are nearly identical to that of nontreated and HU treated 9^KO^ cells (**Fig. 4a**, plots 5&6), highlighting the reproducibility of our methods and providing confidence for fiber quantification throughout.

**Figure 4.**
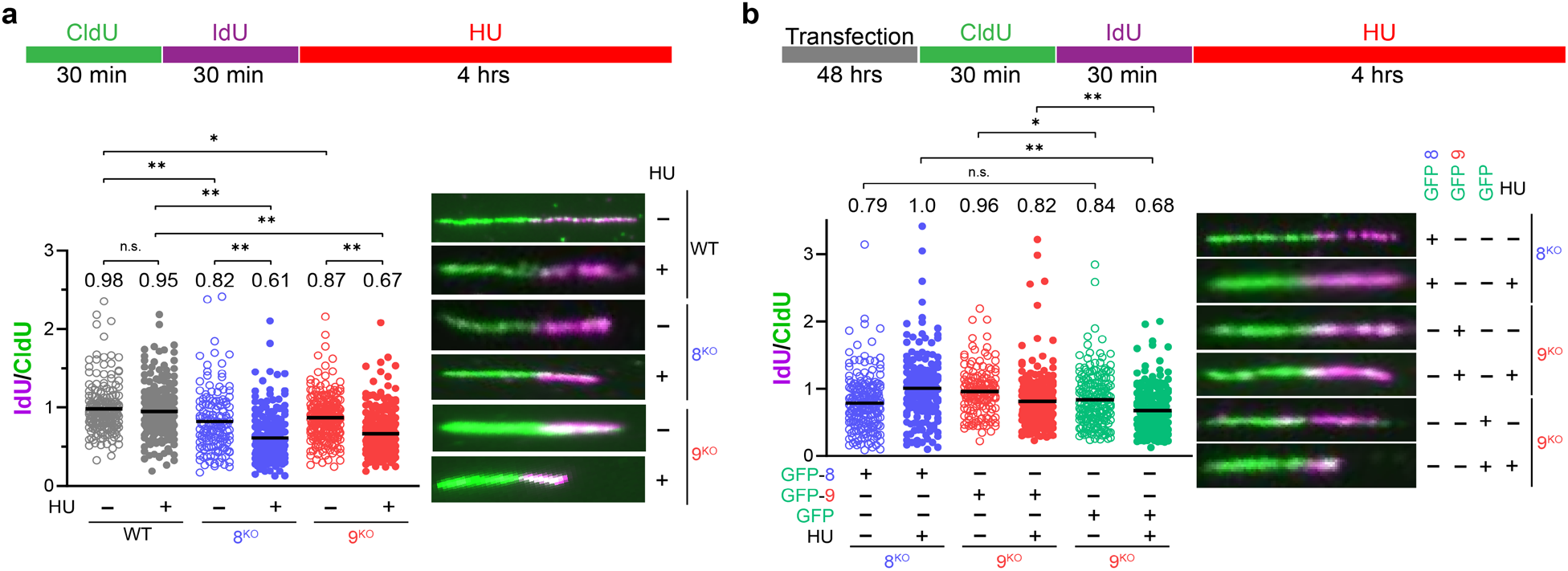
Replication forks are unstable in the absence of MCM8/9. **a** 293T WT (grey), 8^KO^ (blue), or 9^KO^ (red) cells were labeled with CldU followed by IdU for the indicated time intervals (○) or followed by 2 mM HU for 4 hours (●). **b** 8^KO^ (blue), or 9^KO^ (red) cells were transfected with GFP-MCM8, GFP-MCM9, or GFP alone for 24 hours before sequential incubation of CldU and IdU for the indicated time intervals (○) or followed with 2 mM HU for 4 hours (●). DNA was spread by gravity to measure replication fork stability by quantifying fiber lengths using Image J. IdU and CldU lengths were measured using ImageJ and the corresponding ratios reported at the top of the plots and by a black line embedded in the data. Representative fibers are shown to the right of the dot plots. A Mann-Whitney U test was used to calculate P values (n.s. nonsignificant, *P< 0.05, **P< 0.01).

Several SNF2 enzymes (SMARCAL1, HLTF, ZRANB3) catalyze replication fork reversal upon stalling. To examine whether MCM8/9 actively stabilizes forks reversed by these enzymes, SMARCAL1 and HLTF were knocked down separately by siRNA transfection in 8^KO^ or 9^KO^ cells, and fork stability was determined by DNA fiber analysis. siSMARCAL1 rescued the minor decrease mean IdU/CldU ratio values in both nontreated 8^KO^ and 9^KO^ to WT levels (**Fig. 5a**, compare plots with open circles with **Fig. 4a** open circles). This rescue in mean IdU/CldU ratio values was also observed after treatment with 2 mM HU (**Fig. 5a**, compare plots with filled circles with **Fig. 4a** filled circles). These data suggest that, in the absence of MCM8/9, replication fork stability is reduced and prone to degradation following more prevalent reversal by SNF2 fork remodeling enzymes.

**Figure 5.**
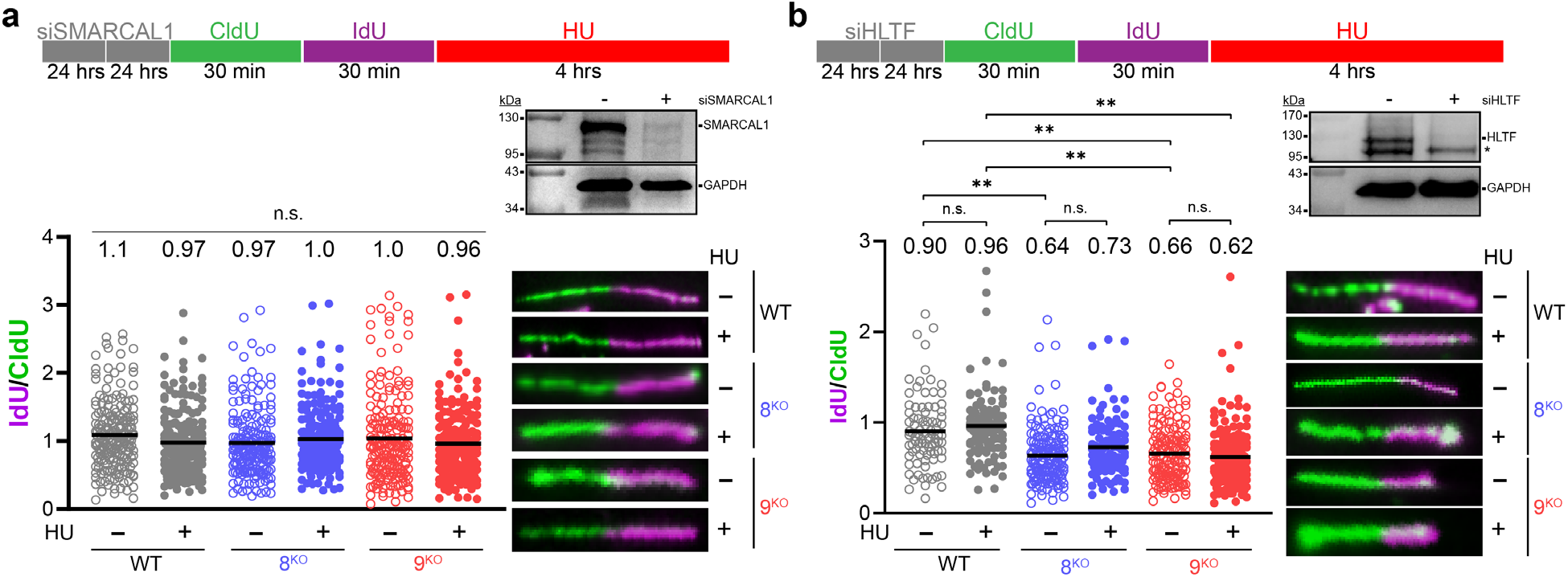
MCM8/9 function in the SMARCAL1-mediated fork reversal mechanism. 293T WT (grey), 8^KO^ (blue), or 9^KO^ (red) cells were transfected with siRNA twice for 24 hours each to knockdown **a** SMARCAL1 or **b** HLTF and verified by western blot. * indicates a nonspecific band on the HLTF blot. Cells were then sequentially incubated with CldU and IdU for the indicated time intervals (○) or followed by 2 mM HU for 4 hours (●). DNA was spread by gravity to measure replication fork stability by quantifying fiber lengths using Image J. IdU and CldU lengths were measured using ImageJ and the corresponding ratios reported at the top of the plots and by a black line embedded in the data. Representative fibers are shown to the right of the dot plots. A Mann-Whitney U test was used to calculate P values (n.s. nonsignificant, **P< 0.01).

Conversely, addition of siHLTF to both nontreated 8^KO^ and 9^KO^ cells reduced IdU/CldU ratios to levels analogous to that observed for 2 mM HU treated conditions (**Fig. 5b**, compare open and filled circles), and unlike for siSMARCAL1, fiber ratios were not rescued. No significant change in fork stability was observed in the WT cells. The reduction in replication fork stability following siHLTF in both nontreated and treated conditions suggests that MCM8/9 function in a complementary but non-overlapping replication fork protection pathway. Indeed, HLTF has been reported to protect replication forks via alternative mechanisms.^36^ Instead, MCM8/9 primarily functions to stabilize reversed replication forks contained within the SMARCAL1 axis.

### MCM8/9 counteracts and restricts BRCA1’s role in fork protection

During HR, BRCA1 supports end resection to generate a 3’ overhang, recruits BRCA2 to the site of damage, and aids in loading (w/BRCA2) of Rad51 onto single-stranded DNA^8,37^, resulting in protection of nascent DNA strands from degradation by the nuclease MRE11.^38^ Fork stability can be restored in BRCA1 deficient cells through inhibition of any one of the SNF2 fork reversal enzymes: SMARCAL1, HLTF, or ZRANB3^34^ or nucleases: MRE11, DNA2, MUS81, and SLX4-ERCC1.^39^ Similarly, we wondered if knockdown of BRCA1 could restore fork stability in MCM8/9 knockout (KO) cells.

DNA fibers were used to examine the role of MCM8/9 in stabilizing HU stalled forks in the absence of BRCA1. As expected, siBRCA1 reduced IdU/CldU ratios in HU stalled WT 293T cells (**Fig. 6a**, plots 1 & 2). However, fork stability was restored when BRCA1 was knocked down in 8^KO^ or 9^KO^ cell lines (**Fig. 6a**, plots 2 vs. 4 or 6). Thus, it appears that stabilization of stalled replication forks is compromised in the absence of either BRCA1 or MCM8/9 but is restored when both are absent. This may be explained by the inability to form reversed (unprotected) forks when both BRCA1 and MCM8/9 are absent. These results emphasize a non-redundant role for MCM8/9 and BRCA1 in maintaining replication fork stability, placing them in the same pathway.

**Figure 6.**
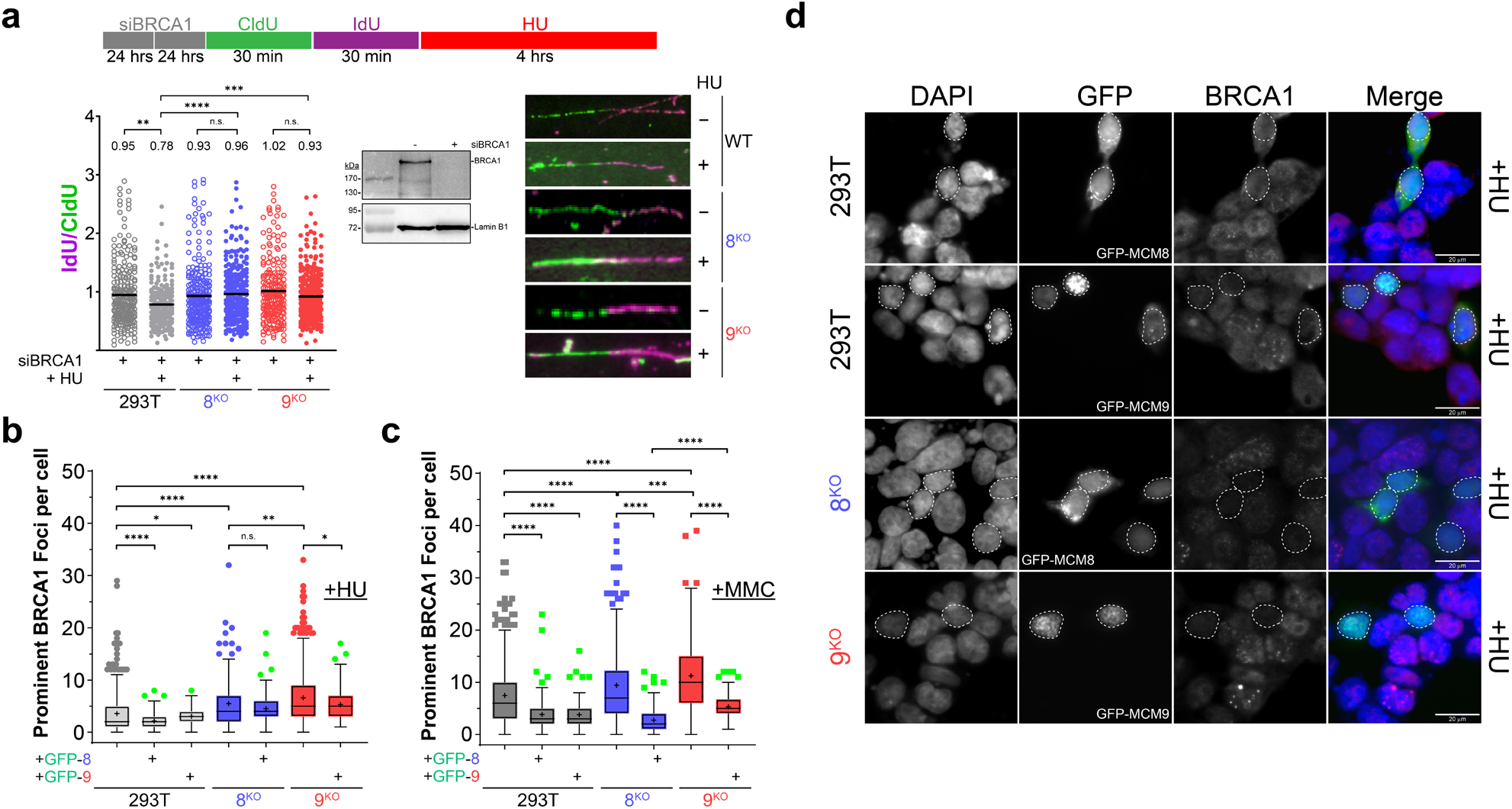
MCM8/9 functions with BRCA1 to maintain replication fork integrity. **a** 293T WT (grey), 8^KO^ (blue), or 9^KO^ (red) cells were transfected with siRNA twice for 24 hours each to knockdown BRCA1 and verified by western blot. Cells were then sequentially incubated with CldU and IdU for the indicated time intervals (○) or followed by 2 mM HU for 4 hours (●). DNA was spread by gravity to measure replication fork stability by quantifying fiber lengths using Image J. Id<u>U</u> and CldU lengths were measured using ImageJ and the corresponding ratios reported at the top of the plots and by a black line embedded in the data. A Mann-Whitney U test was used to calculate P values (n.s. nonsignificant, *P< 0.05, ***P< 0.001, ***P< 0.001, ****P< 0.0001). Representative fibers are shown to the right of the dot plots. GFP-MCM8 or GFP-MCM9 (green, ● or ■) was transfected into each cell line and treated with **b** 2 mM HU for 4 hours (●) or **c** 3 μM MMC for 6 hours (■). Prominent BRCA1 foci were quantified per cell using Image J and plotted as a Tukey box and whiskers with + representing the mean. Representative microscopy images of cells treated with MMC are shown in Supplementary Figure 2. Student’s t-test was used to calculate P values (n.s. nonsignificant, *P< 0.05, **P< 0.01, ***P< 0.001, ****P< 0.0001). d Shows representative microscopy images of cells treated with HU. Dashed white outlines of nuclei indicate cells successfully transfected and show a decrease in BRCA1 signal and foci.

To investigate the dependence and temporal recruitment of BRCA1 in relation to MCM8/9, cells were transfected with GFP-MCM8 or GFP-MCM9 and BRCA1 foci were counted in NT and HU or MMC treated cells (**Fig. 6b-d** and **Supplementary Fig. 2**) Interestingly, the presence or overexpression of MCM8 or MCM9 repressed the formation of BRCA1 foci in treated cells. This was evident in HU treated cells with significant reduction in BRCA1 foci in all cell lines except 8^KO^ which was reduced but just outside the 95% confidence level. The effect was even more pronounced in MMC treated cells, with a significant reduction in BRCA1 foci in all cell lines. This effect was also visually apparent in cells transfected with GFP-MCM8 or GFP-MCM9, where there was a void in BRCA1 foci and signal, unlike in untransfected cells (**Fig. 6d** and **Supplementary Fig. 2**). Therefore, MCM8/9 likely acts to antagonize BRCA1-mediated fork processing activity to maintain fork stability during normal replisome progression, unless forks become severely compromised and stalled.

### The BRCv motif in MCM9 and not helicase activity is necessary to maintain fork stability

Previously, we had characterized a BRC variant motif (BRCv) within the C-terminus of MCM9 (**Fig. 7a**) that interacted with and recruited RAD51 to sites of MMC induced DNA damage.^30^ Therefore, we sought to investigate the role of this MCM9-BRCv motif in maintaining fork stability after HU treatment using DNA fiber analysis. Interestingly, in the absence of HU, fork stability is restored in 9^KO^ cells when MCM9(BRCv^-^) is transfected, implying that the MCM8/9 complex on its own provides some stabilizing context to active replisomes, possibly through its helicase activity (**Fig. 7b**, plots 1 & 3). This increase in fork stability was equivalent to that of adding WT MCM9 back in 9KO cells (compare with **Fig. 4b**, plot 3). When treated with HU and transfected with MCM9(BRCv^-^), fork stability is reduced to basal levels (**Fig. 7b**, plots 2 & 4) and lower than that for adding WT MCM9 (**Fig. 4b**, plot 4), suggesting that an interaction and recruitment of Rad51 is required to provide stabilization to more persistent HU reversed forks through the BRCv motif of MCM9.

**Figure 7.**
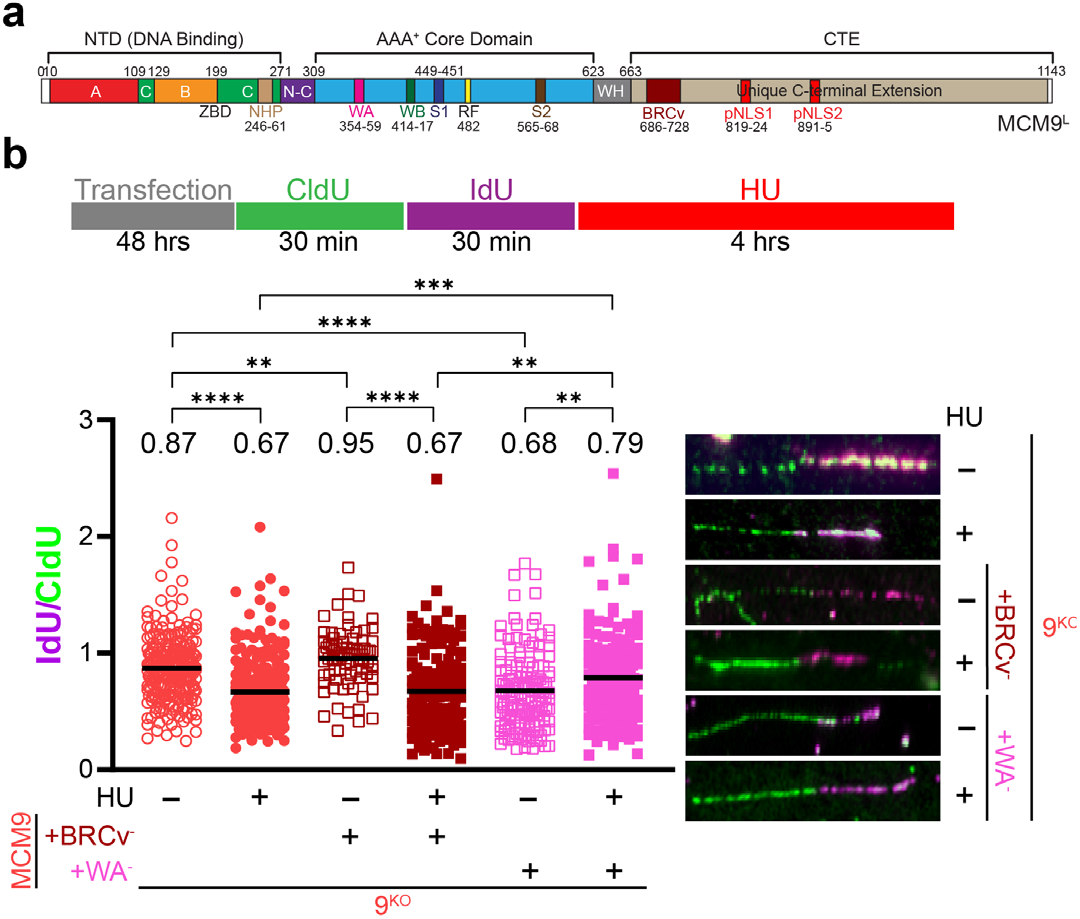
MCM9-BRCv motif is influential in stabilizing HU stalled forks. **a** Schematic of MCM9 showing domains and individual motifs. **b** 9^KO^ cells (○) were transfected pEGFPC2-MCM9(FR687/8AA) (BRCv^-^) (□, brown) or pEGFPC2-MCM9(K358A) (WA^-^) (□, pink) for 48 hours prior. Cells were then sequentially incubated with CldU and IdU for the indicated time intervals or followed by 2 mM HU for 4 hours (● or ■). DNA was spread by gravity to measure replication fork stability by quantifying fiber lengths using Image J. Id<u>U</u> and CldU lengths were measured using ImageJ and the corresponding ratios reported at the top of the plots and by a black line embedded in the data. Representative fibers are shown to the right of the dot plots. A Mann-Whitney test was used to calculate P values (n.s. nonsignificant, **P< 0.01, ***P< 0.001, ****P< 0.0001).

Based on this separation of function mutation for MCM9(BRCv^-^) that is distinct for normal fork progression compared to more persistent stalls, we sought to further investigate the helicase activity of MCM9 by mutating the Walker A site (K358A) and examining its effect on fork stability. Transfection of MCM9(K358A) into nontreated 9^KO^ cells did not rescue fork stability (**Fig. 7b**, plots 1 & 5) unlike that for MCM9(BRCv^-^) above (**Fig. 7b**, plots 3 & 5). Instead, transfection of MCM9(K358A) rescued fork stability only in the presence of HU (**Fig. 7b**, plots 2 & 6), suggesting that recruitment of Rad51 by MCM9, through the BRCv motif, and not direct helicase activity on its own is necessary to stabilized persistently stalled forks.

### MCM8/9 stabilizes stalled forks and protect from nucleolytic degradation

Several nucleases (MRE11, EXO1, and DNA2) have reported activities in processing reversed replication forks to initiate fork recovery and restart.^38,40,41^ However, when excessive fork stalling occurs or when fork protectors are deficient or absent, dysregulated nucleolytic fork degradation by these nucleases is hypothesized to be a source of genomic instability. Based on these previous observations, we hypothesized that the fork instability in both 8^KO^ and 9^KO^ cells was a result of aberrant or excessive nucleolytic degradation and inhibition or knockdown of these nucleases might restore fork stability.

After knockdown of MRE11, EXO1, or DNA2 by siRNA in both 8^KO^ and 9^KO^ cells, fork stability was examined by DNA fiber analysis (**Fig. 8**). Knockdown of MRE11 in both 8^KO^ or 9^KO^ cell lines restored the minor fork instability in nontreated cells to WT levels (**Fig. 8a**, plots with open circles and compare with **Fig. 4**). Interestingly, there was a minor but significant decrease in fork stability in WT cells treated with 2 mM HU (**Fig. 8a**, plots 1 & 2), which highlights the activity of multiple nucleases involved in reversed fork degradation. The addition of 10 μM Mirin (a MRE11 inhibitor) also restored replication fork stability in 8^KO^ and 9^KO^ cells treated with HU but not in the nontreated conditions (**Fig. 8b**, compare open and filled blue, lanes 3 & 4, or red circles, lane 5 & 6) suggesting that alternative forms of fork degradation (*i.e*.EXO1 or CtIP and DNA2) are still active in the absence of HU stress and MCM8/9.

**Figure 8.**
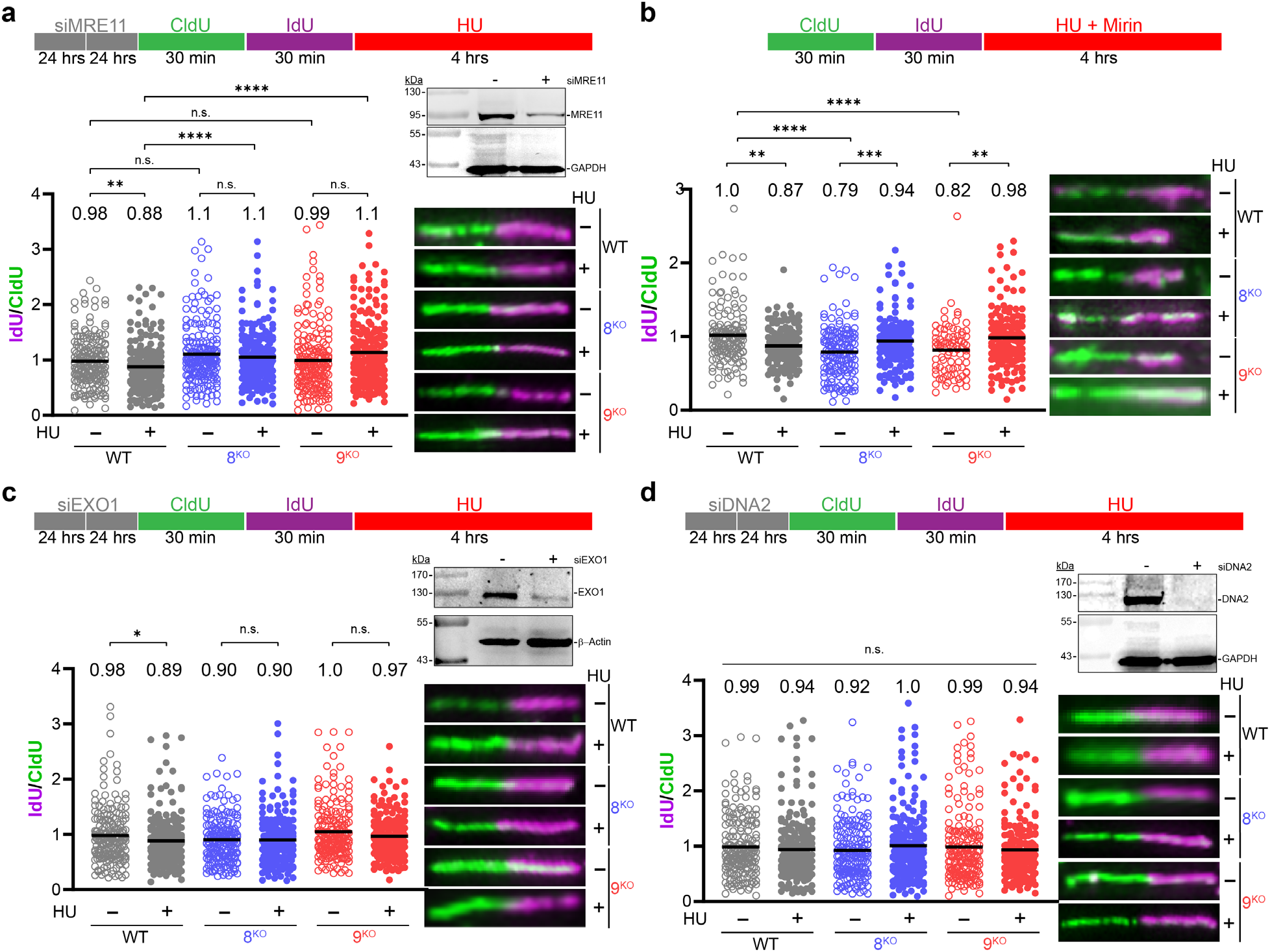
MCM8/9 prevent nucleolytic degradation of replication forks. 293T WT (grey), 8^KO^ (blue), or 9^KO^ (red) cells were transfected with siRNA twice for 24 hours each to knockdown **a** Mre11, **b** treated with 10 μM mirin, or knockdown of **c** Exo1 or **d** Dna2. siRNA knockdown was verified by western blot. Cells were then sequentially incubated with CldU and IdU for the indicated time intervals (○) or followed by 2 mM HU for 4 hours (●). DNA was spread by gravity to measure replication fork stability by quantifying fiber lengths using Image J. Id<u>U</u> and CldU lengths were measured using ImageJ and the corresponding ratios reported at the top of the plots and by a black line embedded in the data. Representative fibers are shown to the right of the dot plots. A Mann-Whitney U test was used to calculate P values (n.s. nonsignificant, *P< 0.05, **P< 0.01, ***P< 0.001, ****P< 0.0001).

Knockdown of EXO1 (**Fig. 8c**) showed a similar trend in fork stability restoration comparable to that observed for siMRE11 (**Fig. 8a**). This is consistent with the two nucleases working in concert to process stalled or reversed replication forks. Lastly, knockdown of DNA2 restored replication fork stability across all conditions examined including WT (**Fig. 8d**), implicating DNA2 as a backup nuclease that processes a specific subset of stalled forks. Overall, this suggests that MCM8/9 has a general protective role in preventing nucleolytic degradation of transiently and more severely stalled replication forks.

### MUS81 robustly cleaves stalled forks in the absence of MCM8/9

We next wanted to address if MCM8/9 are directly involved in replication fork restart. In this experiment, we treated cells with CldU for 30 minutes followed by co -treatment with 2 mM HU with 10 μM Mirin (to prevent nucleolytic degradation of CldU tracts) followed by release from HU and incubation in IdU for 30 minutes to allow stalled replication forks to restart. Both 8^KO^ and 9^KO^ cells did not efficiently restart replication forks compared to WT (**Fig. 9a**, filled plain circles, plots 1 vs. 2 & 4). Interestingly, transfection of GFP-tagged MCM8 or 9 into their respective KO cells also did not efficiently restore replication fork restart (**Fig. 9a**, green outlined circles, plots 3 & 5 vs. 2), suggesting that the MCM8/9 complex is not directly involved in fork restart activities or it promotes alternative mechanisms (such as HR-mediated restart) of replication fork restart. Transfection of the GFP only control in 9^KO^ cells did not allow for efficient restart, as expected (**Fig. 9a**, green filled circles, plot 6).

**Figure 9.**
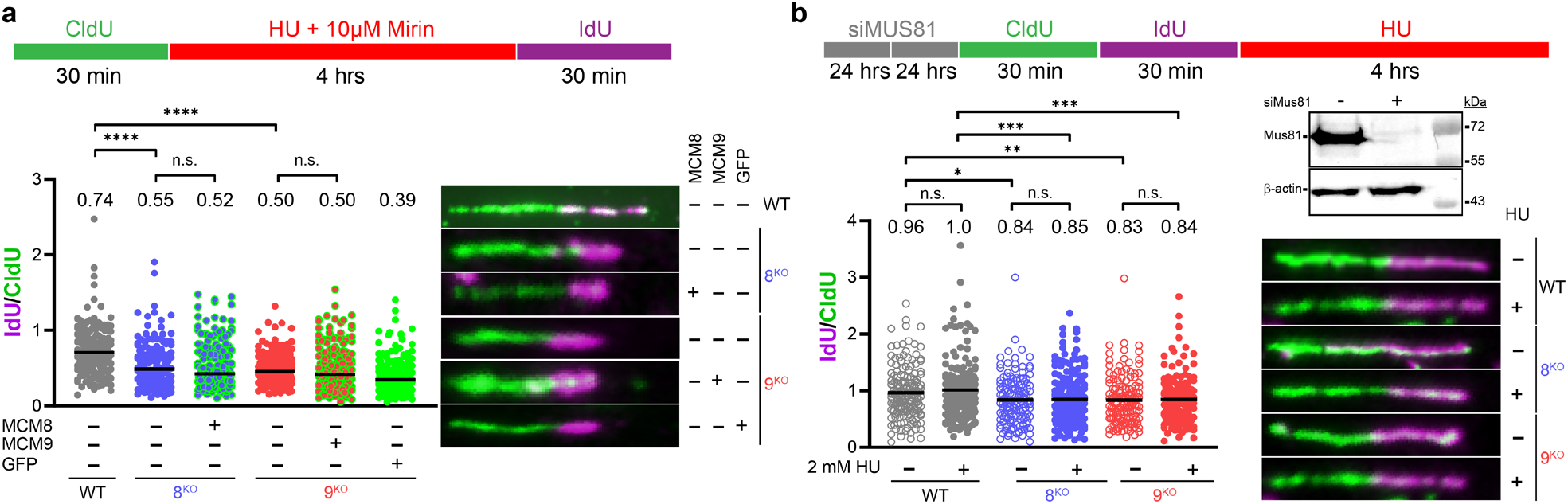
Forks stalled upon MCM8/9 loss are processed by MUS81. **a** 293T WT (grey), 8^KO^ (blue), or 9^KO^ (red) cells were treated with 2 mM HU and 10 μM mirin for 4 hours to monitor fork restart as flanked by CldU and IdU labelling for indicated time intervals. Cells were transfected with MCM8-GFP (green outline, 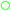), MCM9-GFP (green outline, 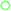), or GFP alone (solid green, 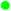) as indicated for 24 hours before initiating fork stalling and restart. **b** Cells were transfected with siRNA twice for 24 hours each to knockdown MUS81 and then sequentially incubated with CldU and IdU for the indicated time intervals (○) or followed by 2 mM HU for 4 hours (●). DNA was spread by gravity to measure replication fork stability by quantifying fiber lengths using Image J. Id<u>U</u> and CldU lengths were measured using ImageJ and the corresponding ratios reported at the top of the plots and by a black line embedded in the data. Representative fibers are shown to the right of the dot plots. A Mann-Whitney U test was used to calculate P values (n.s. nonsignificant, *P< 0.05, **P< 0.01, ***P< 0.001, ****P< 0.0001).

Persistent replication fork stalling and inefficient restart often leads to MUS81-mediated cleavage to initiate HR-mediated repair or fork restart. To investigate whether forks stalled in MCM8/9 KO cells are cleaved by MUS81, we knocked down MUS81 using siRNA and examined fork stability by DNA fiber analysis. Knockdown of MUS81 did not restore replication fork stability in either nontreated 8^KO^ or 9^KO^ cells (**Fig. 9b**, blue or red open circles, plots 3 &4). However, siMUS81 in 8^KO^ and 9^KO^ treated with 2 mM HU restored replication fork stability to levels observed in the nontreated conditions (**Fig. 9b**, compare blue and red open and closed circles, plots 3 vs. 4 and 5 vs. 6). It is known that replication forks are minimally processed by nucleases such as MRE11 prior to generating substrates amenable to MUS81 cleavage.^38^ Our data support a model in which fork stability is not completely restored in nontreated cells depleted for MUS81, as nucleases are still present to minimally process reversed forks. However, when forks are persistently stalled with HU, fork stability is restored to basal levels in 8^KO^ or 9^KO^ cells when MUS81 is knocked down (**Fig. 9b**, compare closed circles, plots 4 & 6 with those in **Fig. 4a**, plots 4 & 6).

## Discussion

Mutations in MCM8 and MCM9 have been clearly linked with infertility and primary ovarian insufficiency^14,15^ as well as predispositions to a variety of cancers.^42,43^ The MCM8/9 complex has been primarily correlated with a role in DSB repair from damage induced by MMC, cis-PT, or IR contributing to HR^19,21,24,31^, however, MCM8/9 has also been detected directly at replication forks.^27,28^ This prompted us to investigate whether MCM8/9 also participates during active replication to either protect, promote, or process stalled replication forks. Our results are consistent with a fork protection role for MCM8/9 that occurs during active replication fork progression in the absence of any exogenous damage, responding to transient impediments to replication, as well as during more severe replisome stalling induced by HU (**Fig. 10**).

**Figure 10.**
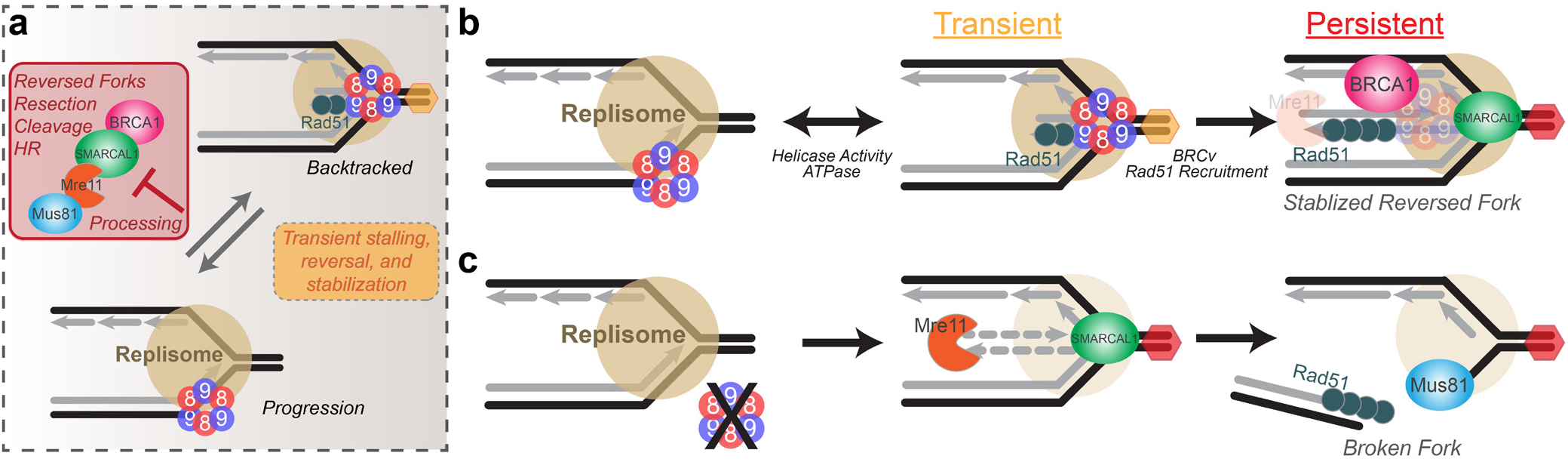
A model for MCM8/9 aiding in fork progression with transient or persistent obstacles. **a** MCM8 and 9 participate within the replisome to overcome transient blocks to fork progression for protection and to maintain genomic integrity. **b** Should the replisome encounter persistent blocks, MCM8/9 works in the SMARCAL1 fork reversal pathway to recruit BRCA1 and Rad51 to stabilized stalled forks for restart. c In the absence of MCM8/9 and endogenous stressors, transiently reversed forks become more persistent and are processed through a SMARCAL1/MRE11/MUS81 pathway, resulting in increased in DBSs.

We can now show that MCM8/9 normally aids in maintaining fork progression and that their absence results in severe fork instability leading to DSBs induced by MUS81. Previously, targeted depletion of the MCM2 subunit of the MCM2-7 replication fork helicase complex resulted in continued replication and synthesis by MCM8/9, albeit at a significantly slower rate^27^, consistent with our finding. MCM8 and MCM9 have been detected at higher abundances than even MCM2-7 (due to loading at dormant origins) or any other helicase at replication forks from coupled immunoprecipitation mass spectrometry studies.^28^ Therefore, MCM8/9 facilitate replisome progression, and when they are absent, genome stability suffers as indicated by significantly more γH2A.X foci and staining, even in the absence of any exogenous stressors. Although γH2A.X is commonly utilized as a marker for DSBs, it can also mark persistently blocked forks or single strand breaks directed by ATM.^44^ In fact, our neutral comet analysis did not show significant DSBs in nontreated 8^KO^ or 9^KO^ cells, consistent with single breaks or significant stalling. Combined, these results suggest that MCM8/9 is present and active within replisomes to aid in fork progression through challenging genomic stretches that may result in transient fork stalling/reversal processes.

Upon more severe fork stalling initiated by HU (or MMC), effects of MCM8 or MCM9 KO on genome stability become more evident. MCM8 and MCM9 form nuclear foci when cells are treated with a variety of DNA damage agents, and in their absence, more DSBs are detected by longer comet tail moments. To investigate the consequences to fork stability upon KO of MCM8 and MCM9, we utilized a suite of DNA fiber assays to specifically probe fork reversal, protection, and resection. DNA fiber analysis shows that knockdown of SMARCAL1 and not HLTF restores fork stability, overall implicating increased SMARCAL1 fork reversal activity when MCM8 or MCM9 are deficient. Resection of stalled forks is complicated by several nucleases acting with overlapping specificities and cooperativities to degrade a spectrum of reversed fork structures. Knockdown of MRE11 appears to have the greatest effect in restoring DNA fiber lengths in 8^KO^ and 9^KO^ cell lines, which was also corroborated with separate treatment with Mirin. However, siEXO1 also restored DNA fiber lengths, similar to that of siMRE11. siDNA2 was interesting in that in addition to restoring stability in 8^KO^ and 9^KO^ cells, it also completely restored the minimal sensitivity seen in WT cells. In our studies, the defects seen with HU stalled forks in MCM8 or 9 deficient cells are linked to processes prior to fork resection and adding back either MCM8 or 9 did not rescue fork restart. This result is slightly different than that shown previously where MCM8/9 aided more directly in MRN resection processes of severely reversed forks.^21^ Even so, in the absence of MCM8 or MCM9, stalled forks become extremely unstable, are actively reversed by SMARCAL1, and then resected by a combination of coordinating nucleases to process all types of intermediates.

As both MCM8^KO^ and MCM9^KO^ cell lines have increased γH2A.X foci in the absence and presence of exogenous agents, it is likely there is a spectrum of DNA intermediates with single strand gaps, stalled replisomes, and various reversed fork structures that require MCM8/9 response. Once restart processes fail, those intermediates become targets of MUS81 cleavage. In fact, knockdown of MUS81 restored some fork stability under HU stalled conditions but not completely, highlighting again the competing roles of other nucleases and sub-pathways in this process. One of the hallmarks of MCM8 or MCM9 patient deficient cells was extreme sensitivities to MMC and the formation of broken, fused, and radial chromosomes^14,15^, consistent with DNA end-joining processes occurring after more rampant fork cleavage by MUS81 and defects in HR.

Our evidence places MCM8/9 within or around the replisome actively responding to both transient and persistent stalling events to protect forks before significant processing can take place. An interaction with RAD51 through the MCM9 BRCv motif, in particular, is influential in this dynamic protection process and likely stabilizes a subset of stalled forks that are not significantly reversed.^30^ Even transfection of a catalytically inactive MCM9 stabilizes forks to a greater level than that of a BRCv^-^ mutant during HU stalling, highlighting the importance of the BRCv motif to recruit Rad51 over that of the any associated ATPase activity utilized for fork progression. MCM8/9 antagonizes the effects of BRCA1 localization, which itself acts to stabilize stalled and reversed forks slightly further downstream to aid in their restart.^45^ Although BRCA1 and BRCA2 are generally assumed to play similar but temporal roles in fork stability, to sequentially recruit Rad51, emerging data suggests they are affected differently by MUS81.^38,39^ While depletion of MUS81 confers fork protection in BRCA2^-/-^ cells through a break induced replication (BIR) pathway, it does not in BRCA1^-/-^ deficient cells, highlighting a divergence in repair pathways, where BRCA1 can more adequately protect reversed forks from cleavage. Interestingly, more complete fork restoration required the elimination of both MCM8/9 and BRCA1, suggesting that fork reversal may not be possible in this situation as no suitable nuclease substrates are formed. In that case, an alternative pathway such as MUS81 cleavage and HR may be utilized. Therefore, MCM8/9 and BRCA1 appear to have nonredundant roles in maintaining fork stability, where MCM8/9 are acting prior to BRCA1 recruitment but are exclusive to one another.

Altogether, MCM8/9 mediates early fork stalling and stabilization processes that aid in active replication processes, but they also work to recruit downstream reversed fork protection proteins, including Rad51 and BRCA1 when forks are more severely stalled. The ATPase activity of MCM8/9 is utilized for normal fork progression, possibly to restrict the formation of transiently reversed fork structures that can be recognized by stabilizers. However, upon severe stalling, MCM9 recruits RAD51 to initiate fork reversal and restart processes. In the future, it will be interesting to determine how MCM8/9 is incorporated within the replisome, better understand its DNA substrate specificity used modulate reversed forks and interrogate the role of MCM9 recruited Rad51 in initiating downstream repair processes.

## Materials and Methods

### Cell Culture

8^KO^ and 9^KO^ in parental 293T cells were created using CRISPR/Cas9 technology and confirmed knockout by DNA sequencing and mitomycin C (MMC) sensitivity assays.^30^ All cells were cultured at 37 °C with 5% CO_2_ in Dulbecco’s modified eagle’s medium (DMEM) (Corning Cellgro) supplemented with fetal bovine serum (FBS) (Atlanta Biologicals) at a 10% working concentration. Plasmid transfections (pEGFPc2-MCM8, pEGFPc2-MCM9, pEGFPc2-MCM9(FR687/8AA) were carried out using LPEI (ThermoFisher) as described.^30^ The MCM8 or MCM9 gene was cloned into the pIRES2-EGFP using traditional restriction site cloning, *BglII/BamHI* and *XhoI/XmaI*, respectively. pEGFPc2-MCM9(K358A) was created by a modified QuikChange protocol and screened with a novel inserted restriction enzyme, *SmaI* (NENB). siRNAs were obtained form Dharmacon for

siSMARCAL1 (5’-GCUUUGACCUUCUUAGCAAUU),
siHLTF (5’-GGUGCUUUGGCCUAUAUCAUU),
siBRCA1 (5’-CUAGAAAUCUGUUGCUAUG),
siMre11 (5’-ACAGGAGAAGAGAUCAACU),
siDNA2 (5’-GUAACUUGUUUAUUAGACAUU), and
siMus81 (5’-CAGCCCUGGUGGAUCGAUAdTdT)

or from Sigma for universal negative control #1 (SIC001) and were transfected using TransIT-X2 (Mirus) in Opti-MEM media following manufacturer’s recommendation. Cells were transfected twice for 24 hours with the indicated siRNA before treatments. Cells were then treated with either 2 mM HU (Acros) for 4 hours, 3 μM MMC (ThermoFisher) for 6 hours, or 10 μM mirin (Sigma) for 6 hours by adding agent directly to the media.

### Western Blotting

Harvested cells were lysed in RIPA buffer (50 mM Tris pH 7.5, 150 mM NaCl, 0.1% SDS, 0.05% Triton, 10 mM DTT, 0.5 μM EDTA) and sonicated on ice. Protein content was quantified by BCA Assay (Boster Bio, AR01466) and stored at −20 °C. 30 μg of lysed protein was thawed on ice, electrophoresed on 8% or 10% acrylamide SDS-PAGE gel, and transferred onto PVDF or nitrocellulose in transfer buffer (25 mM Tris-HCl [pH 7.6], 192 mM glycine, 20% MeOH, 0.0375% SDS). The membrane was cut and blocked overnight in 5% powdered milk in 1xTBST at 4 °C, rocking. Following a wash with 1x TBST (used for all washes), membranes were incubated with their respective primary antibody [α-Smarcal1 (Bethyl Laboratories, A301-616A), 1:100; α-HLTF (Bethyl Laboratories, A300-640A), 1:1000; α-BRCA1 (Santa Cruz, sc-6954), 1:250; α-MRE11 (Proteintech, 10744-1-AP), 1:500; α-DNA2 (Invitrogen, PA5-68167), 1:100; α-EXO1 (Bethyl Laboratories, A302-640A), 1:2000; α-MUS81 (Abcam, ab14387), 1:1000; α-GAPDH (Pierce, MA5-15738), 1:20,000, α-Lamin B1 (Proteintech, 12987-1-AP), 1:5,000, α-β-actin (Abcam, ab82227), 1:10,000 for two hours, rocking, at room temperature. The membranes were washed three times and incubated with secondary antibodies (goat anti-rabbit HRP Novex, A16096) (goat anti-mouse HRP, Novex, A16072), ranging from 1:1000-10,000, for one hour, rocking, at room temperature. Three more washes were performed before addition of luminol reagents (Santa Cruz) and/or imaging with a Typhoon FLA9000 or ImageQuant LAS 4000 (Cytiva, Marlborough, MA) imagers.

### DNA Fibers

DNA fiber assays were performed as described previously with slight optimization modifications.^46,47^ Briefly, cells were treated sequentially with 50 μM IdU (TCI America) and 500 μM CldU (MP Biomedicals) nucleotide analogs for 30 minutes each, with a gentle wash with 1X PBS in between nucleotide incubations, prior to (unless indicated otherwise) treatment with DNA damaging or fork stalling agents. Cells were harvested after 2 washes with 1X PBS, pelleted and stored at −20 °C before spreading. Cells were spread by gravity on silanized microscope slides by mixing 2 μL of cell suspension with an 8 μL drop of DNA fiber lysis buffer (200 mM Tris-Cl, pH 7.5; 50 mM EDTA; and 0.5% SDS). Drops were allowed to dehydrate for 10-20 min prior to spreading. Fiber spreads were then allowed to dry completely and were fixed to the slide by incubating in a 1:3 solution of methanol:acetic acid for 10 minutes before storage overnight at −20 °C. Fixed fibers were denatured for 25 minutes in 1 M NaOH solution followed by 2-3 washes in 1X PBS. Fibers were blocked for 30 minutes in fiber blocking buffer consisting of 1X PBS, 5% bovine serum albumin (BSA), and 0.1% Tween. Fibers were then incubated sequentially in humidified chambers with mouse (BD Bioscience, BD-347580) (1:50) and rat (Abcam, ab6325) (1:400) primary anti-BrdU antibodies in fiber blocking buffer for 1 hour each with 2-3 washes in 1X PBS with 0.1% Tween between incubations. Fibers were simultaneously incubated in α-mouse-Cy3-conjugated (Abcam, 97035) and α-rat 488-conjugated (Abcam, 150157) secondary antibodies (1:400) in fiber blocking buffer for 1 hour. Slides were washed 2-3 times in 1X PBS with 0.1% Tween followed by mounting in mounting media consisting of 0.5X PBS, 25 mg/mL 1,4-Diazabicyclo[2.2.2]octane (DABCO), 1 mM ascorbic acid, and 90% glycerol. Mounted slides were sealed with clear polish. Fibers were then imaged on an Olympus IX-81 epifluorescence microscope with a 60X oil immersion objective and analyzed using Cell Sens Dimension 2 software. 100 or more fiber lengths were measured with ImageJ software (Rasband 1997-2016, 17 October 2015) to calculate IdU/CldU ratio values. For replication rate, slope values in μm m^-1^ were converted to bp s^-1^ using the known base pair distance (3.4 Å bp^-1^) as the conversion factor. Scatter plots were created using GraphPad Prism (v.9.2) and a Mann-Whitney U test was conducted to analyze statistical significances unless indicated otherwise.

### FACS Analysis

293T, 8^KO^, or 9^KO^ cells were synchronized at the beginning of S-phase using a double thymidine block. Adherent cells were grown to 40% confluency in 10 cm^2^ dishes with DMEM/10% FCS supplemented with 10 mM thymidine (TCI America) and cultured at 37 °C with 5% CO_2_. After 18 hours, the media was aspirated, and cells were washed three times with 10 mL pre-warmed PBS. Cells were released by the addition of unsupplemented DMEM/10% FCS for 8 hours. Cells were again synchronized into G1/S phase by addition of DMEM 10%/FCS/10 mM Thymidine. After 18 hours, cells were washed 3x with pre-warmed PBS and released into fresh DMEM 10% FCS. Cells were harvested and fixed in 70% ethanol at indicated timepoints and stored at 4 °C. Cell pellets were stained using PI/RNase Staining Buffer (BD Biosciences, 550825) per manufacturers protocol. The cell cycle profile data was collected on a FACSVerse (BD Biosciences) using the propidium iodide channel. Cell cycle determination was analyzed using forward scatter (FSc) and side scatter (SSc), selecting for unaggregated live cells, graphed using FlowJo (BD Bioscience, v10) and presented using Adobe Illustrator (2021).

### Fluorescence and Immunofluorescence Imaging

Adherent cells on glass coverslips were washed in 1X PBS (2 times), fixed in 4% paraformaldehyde in PBS for 10 minutes and permeabilized with 0.1% Triton X-100 in PBS (PBST) for 15 minutes. Cells were blocked overnight with 5% BSA in PBST at 4 °C. For immunofluorescence, coverslips were incubated with α-γH2A.Xp (Abcam, ab26350) (1:400) or α-BRCA1 (Santa Cruz, sc-6954) (1:50) dilution of primary antibodies in 2.5% BSA in PBST for 1 hour at 37 °C. Cells were washed three times in PBST and incubated with 1:1000 dilution of the α-mouse Alexa647 (ThermoFisher, A-21235) secondary antibody followed and then washed three times with PBST. Cells were mounted in DAPI mountant (Prolong Gold, Thermo Fisher) and sealed with clear polish and imaged under a FV-1000 epifluorescence or FV-3000 confocal laser scanning microscope (Olympus Corp.). Images were processed with vendor included Fluoview (v.4.2b) or CellSens software (dimension 2). γH2A.X or BRCA1 foci from epifluorescence images were automatically counted from individually gated cells using identical thresholds that eliminated background noise using Image J, as described previously.^30^ Foci per cell are presented in a box and whisker plot to identify the upper and lower quartiles, outliers, the median, and the mean. Data was analyzed for any statistically significant differences using a Student’s two-tailed unpaired t-test in GraphPad Prism unless otherwise indicated.

### Neutral Comet Assay

Comet assays were performed with the CometAssay^®^ Electrophoresis System II (Trevigen, 4250-050-ES) following the manufacturer’s protocol. Briefly, cells were harvested in 1X PBS. Cells were diluted in low-melting point agarose to a concentration of 1 x 10^6^ cells/mL and 50 μL of cell solution was spotted on a microscope slide. Slides were placed in the dark at 4 °C for 30-45 minutes to allow the agarose spot to dry. Slides were then immersed in Lysis Solution provided by the manufacturer for 30 minutes, then cooled to 4°C for 60 minutes. Slides were then placed in 1X neutral electrophoresis buffer (50 mM Tris [pH 9.0], 150 mM sodium acetate) for 30 minutes. DNA was then electrophoresed at 21 V for 45 minutes and then the slides were immersed in DNA precipitation buffer (7.5 M Sodium Acetate and 95% ethanol) for 30 minutes at room temperature. Slides were then rinsed in water for 5 minutes followed by 70% ethanol for 5 minutes. Slides were then dried at 37 °C for 10-15 minutes followed by incubation in 1X SYBR Gold DNA stain (ThermoFisher, S11494) for 30 minutes at room temperature. Slides were briefly washed in water to remove excess stain and were allowed to dry completely at 37 °C. Slides were mounted with mounting solution as detailed above and imaged by epifluorescence microscopy. Percent DNA in the comet tails were measured with ImageJ software and tail moments were calculated according to Equations 1–2:

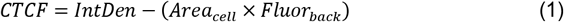

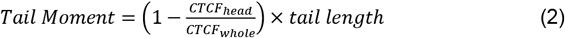

where *CTCF* is the corrected total cell fluorescence for a comet head (*head*) or the whole comet (*whole*), *IntDen* is the integrated density, *Area_cell_* is the area of the selected cell, and *Fluor_back_* is the background mean fluorescence. Scatter plots were created using GraphPad Prism and a Mann-Whitney U test was conducted to analyze statistical significances unless indicated otherwise.

### Quantification and Statistical Analysis

Bars and error bars represent mean and SEM, respectively, of indicated numbers of independent experiments. Statistical analysis was performed by a Student’s two-tailed unpaired t-test or a Mann-Whitney U test in GraphPad Prism, as indicated. Scatter plots show all the individual data points while boxplots show the first quartile, median, third quartile and whiskers which extend to 1.5X of the interquartile range. Outliers are shown.

This is an equation line. Type the equation in the equation editor field, then put the number of the equation in the brackets at right. The equation line is a one-row table, it allows you to both center the equation and have a right-justified reference, as found in most journals.

### Data Availability

Raw data and images are also available upon request. No new codes were generated.

## End Matter

### Author Contributions and Notes

W.C.G. performed and analyzed most of the DNA fiber assays, comet assays, and some of the confocal microscopy. D.R.M. provided the 8^KO^ and 9^KO^ cell lines and performed the FACS analysis. K.N.K did the siBRCA1 fiber assay, transfections, and western blots. R.B. performed transfections, comet assays, and western blots. M.A.T performed the gH2A.X and BRCA1 immunofluorescence assays and quantified the associated foci. M.A.T and W.C.G conceived the project and wrote the manuscript with input from all the authors. M.A.T supervised the project.

The authors declare no conflict of interest.

This article contains supporting information after the references.

## Acknowledgments

The authors wish to thank the Dr. Bernd Zechmann (Center for Microscopy and Imaging, Baylor University, Waco, Texas) and Dr. Michelle Nemec (Molecular Biosciences Center, Baylor University, Waco, Texas) for technical support during this work. We thank Elizabeth Jeffries for cloning the pIRES2-EGFP-MCM8 and pIRES2-EGFP-MCM9 plasmids. This work was supported by Baylor University and a NIH R15 (GM13791 to M.A.T.).

---

**Supplemental Figure S1.**
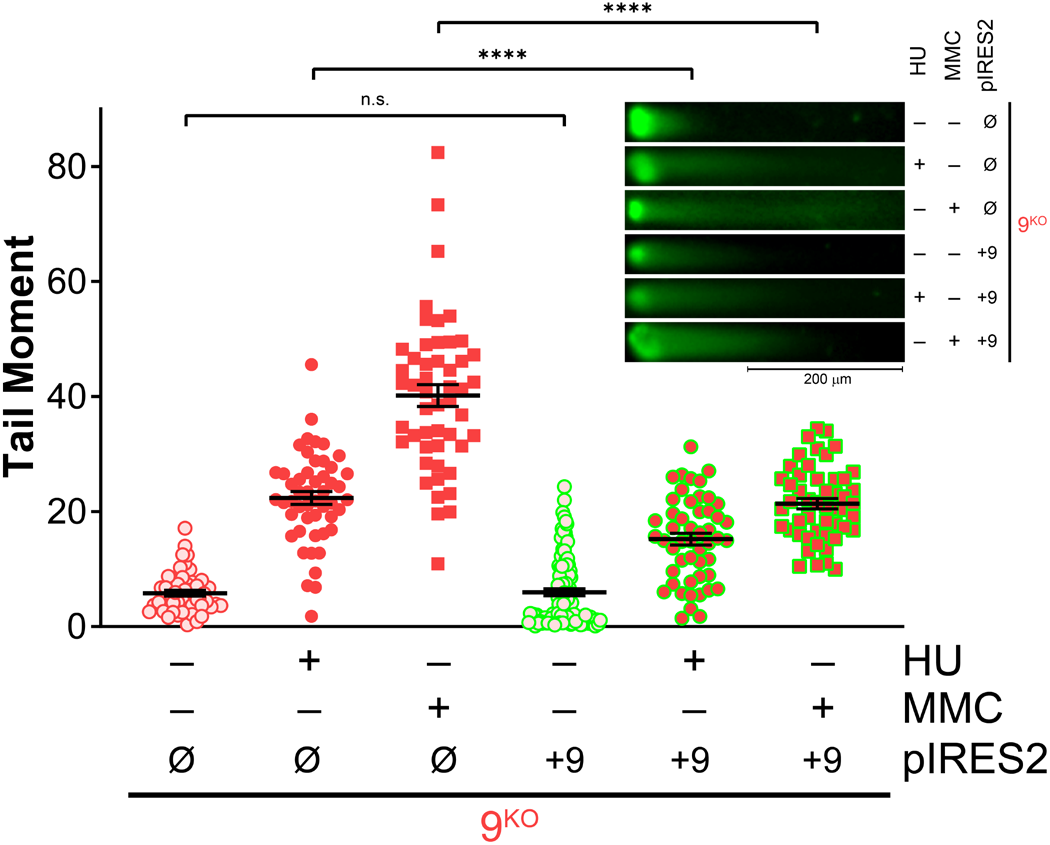
Restoration of Comet tails through addition of MCM9 to 9^KO^. 9^KO^ cells were transfected with pIRES2 empty vector 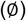 or pIRES2-MCM9 (+9) (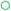, green outline) either nontreated (NT) 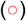 or treated with 2 mM HU for 4 hours (+HU) 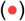 or 3 μM MMC for 6 hours 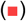. The comet tail moments were calculated with the mean and standard error of the mean reported as black bars and examined for significant differences using a two-tailed unpaired Student’s t-test. (n.s. nonsignificant, ****P< 0.0001). Representative comets are shown inset with the plot with indicated scale bar of 200 μm).

**Supplemental Figure S2.**
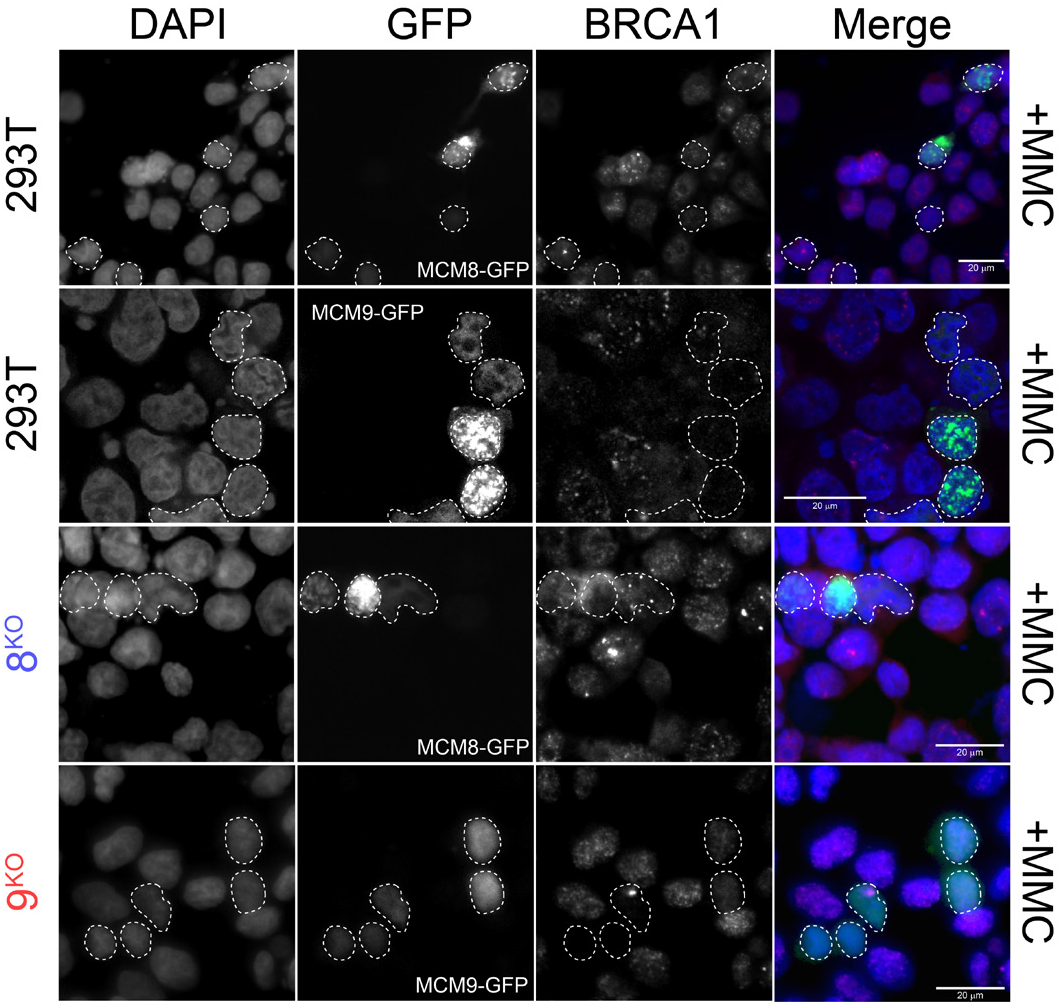
MCM8/9 suppresses BRCA1 foci formation. 293T WT, 8^KO^, or 9^KO^ cells were transfected with MCM8-GFP or MCM9-GFP was transfected and treated with 3 μM MMC for 6 hours. Dashed white outlines of nuclei indicate cells successfully transfected and show a decrease in BRCA1 staining and foci. Quantification of prominent BRCA1 foci was done in Figure 6C. Scale bars represent 20 μm.

